# Epigenetic Control of Spatiotemporal Dynamics of Pancreatic Cancer Cells via Brg1–Rac1 Signaling

**DOI:** 10.64898/2026.05.04.722641

**Authors:** Akihisa Yamamoto, Akihisa Fukuda, Yuichi Fukunaga, Kentaro Hayashi, Hiroshi Seno, Motomu Tanaka

## Abstract

Pancreatic ductal adenocarcinoma (PDA) arises from distinct precursor lesions with different clinical outcomes, yet the mechanisms linking epigenetic regulation to invasive cell behavior remain poorly understood. Here, we investigate how the chromatin remodeler Brg1 influences the dynamic properties of cancer cell migration. Using a biomimetic supported membrane system combined with label-free interferometric imaging, we quantitatively analyze the spatiotemporal dynamics of PDA cells derived from pancreatic intraepithelial neoplasia (PanIN) and intraductal papillary mucinous neoplasms (IPMN). Despite their similar morphology under conventional conditions, PanIN- and IPMN-derived PDA cells exhibit markedly different migration behaviors. PanIN-derived cells migrate faster and display enhanced dynamic remodeling, whereas IPMN-derived cells show persistent elongation with limited displacement. These differences are captured by quantitative analyses of cell trajectories and deformation dynamics. Mechanistically, PanIN-derived PDA cells exhibit elevated Rac1 activity, supporting a model in which a Brg1–Rac1 axis regulates cytoskeletal dynamics and migration behavior. Together, our findings demonstrate that epigenetic regulation is linked to distinct dynamic phenotypes of cancer cells and highlight the importance of quantitative analysis of cell behavior for understanding invasive potential.

## 1. Introduction

Pancreatic ductal adenocarcinoma (PDA) is the sixth leading cause of cancer-related death worldwide, with the overall five-year survival rate of about 10% (Yu *et al*, 2025). Under an increasing number of disease incidence, the interception at an early precancerous stage has become a pushing societal demand (Pedro & Wood, 2025). PDA progresses from well-defined precursor lesions. About 85 % of PDA develop from pancreatic intraepithelial neoplasia (PanIN), and PDA developed from intraductal papillary mucinous neoplasms (IPMNs) are minor. Though both are mucinous, papillary epithelial lesions and exhibit similar driver gene mutations, such as oncogenic mutation of Kras (Fukuda, 2015), the clinical outputs are clearly distinct. PanIN-derived PDA generally has a worse prognosis, higher histologic grade, and more advanced stage at diagnosis compared to IPMN-derived PDA (Habib *et al*, 2024). Previously, Figura et al. found that the loss of chromatin regulator Brg1 (SMARCA4), in combination with oncogenic Kras, causes pancreatic duct cells to form IPMN in mouse models, demonstrating the key role of chromatin remodeling in PDA development (von Figura *et al*, 2014). At present, the design of clinical strategies for PanIN and IPMN encounters several problems, including the difficulties with detection (Pedro & Wood, 2025). For example, typical PanIN are < 5 mm in diameter, which makes them hard to detect by macroscopic imaging methods like CT and MRI (Basturk *et al*, 2015; Kiemen *et al*, 2024).

PanIN-derived cancers are associated with more aggressive clinical features, such as invasive migration into perineural zones and a higher possibility of distant metastasis. On the molecular level, PanIN-derived PDA is associated with enhanced inflammatory pathways and metabolic characteristics, which promote tumor progression. In contrast, IPMN-derived tumors show different genetic pathways that contribute to a slightly more indolent phenotype. However, little is understood how the genetic and epigenetic cues translate into the invasive migration of PDA cells. PanIN and IPMN present genotype-specific morphological phenotypes in precancerous stage. However, Figura et al. reported that morphological phenotypes of PanIN-PDA and IPMN-PDA become hardly distinguishable (von Figura *et al*., 2014). Furthermore, isolated PanIN-PDA and IPMN-PDA cells seeded on Petri dishes show no marked morphological difference. This makes it difficult to link epigenetic remodeling of chromatin by Brg1 to differential invasion and metastasis observed *in vivo*. From a biophysical viewpoint, mesenchymal cell migration is driven by mechanical deformation forces generated by remodeling of focal adhesion contacts and stretch/retraction of actomyosin complexes (Ohta *et al*, 2018; Sackmann & Tanaka, 2021). Therefore, the combination of biomimetic surrogate surfaces and quantitative analyses of cellular phenotypes would enable to highlight how chromatin remodeling by Brg1 modulates mechanical deformation forces and migration of PDA cells in a genotype-dependent manner.

In this study, we investigated spatiotemporal dynamics of IPMN-PDA cells and PanIN-PDA cells by the combination of laminin-functionalized surrogate surfaces and label-free interferometry imaging of living cells. Dynamics of two cell types were analyzed in real space by the analysis of central trajectories and autocorrelation of deformation. Moreover, power spectrum analyses in angular and temporal frequency domains highlight characteristic patterns of differential cellular dynamics. On the protein level, distinct cellular dynamics could be attributed to the differential levels of Rho GTPase activation. These data indicate that Brg1-GEF-Rac1 acts as a regulatory axis connecting chromatin remodeling to divergent migration phenotypes in pancreatic cancer. The integration of quantitative analyses cellular dynamics with molecular signaling mechanisms highlights that critical role of Brg1-dependent Rac1 activation in controlling dynamic cellular phenotypes. The quantitative indices established here could broadly be applied as numerical descriptors assessing the invasive potential of cancer.

## Results

### PDAs derived from distinct precancerous lesions are indistinguishable *in vivo* and *in vitro*

**Figure 1a** presents histopathological images of precancerous IPMN and PanIN lesions of mouse models. The corresponding images of human pathological specimens are shown in **Figure 1b**. In both mouse and human tissues, IPMN lesions are characterized by long and complex papillary projections, while PanIN lesions exhibit epithelium-like phenotypes. However, after progression to cancer, the pathological features of IPMN-PDA and PanIN-PDA in mice become very similar (**Figure 1c**) (von Figura *et al*., 2014). In human, PanIN-PDA and IPMN-PDA are also histologically indistinguishable (**Figure 1d**), which is consistent with previous reports (Basturk *et al*., 2015). When IPMN-PDA and PanIN-PDA cells established from mouse models are seeded on plastic dishes, they are hardly distinguishable, too (**Figure 1e**). In spite of clear differences in molecular and biological characteristics, the morphometric parameters such as projected cell area and aspect ratio show no significant differences (**Figure S1**).

**Figure 1.**
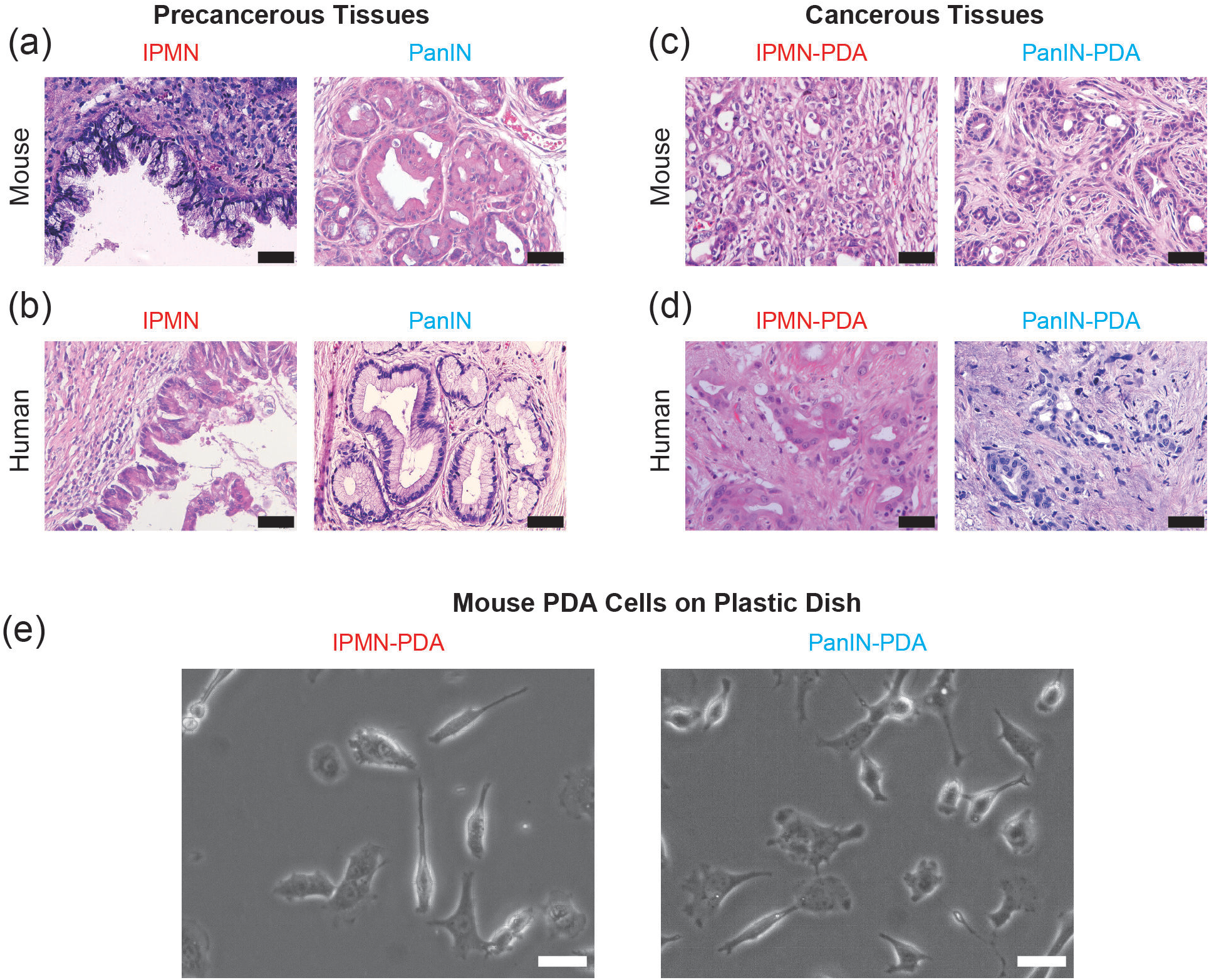
IPMN-derived PDA and PanIN-derived PDA are indistinguishable on tissue and cellular levels. (a) Histopathological images of precancerous lesions (IPMN and PanIN) in mouse models. (b) IPMN and PanIN lesions in human subjects. (c) Histopathological images of IPMN-PDA and PanIN-PDA in mouse models. (d) IPMN-PDA and PanIN-PDA in human subjects. Scale bars (a – d): 50 µm (e) Phase contrast images of mouse IPMN-PDA and PanIN-PDA cells on plastic dishes. Scale bars: 50 µm.

In this study, IPMN-PDA and PanIN-PDA cells are seeded on the membrane-based surrogate surface (**Figure 2a**). The surrogate surface is based on a planar lipid membrane deposited on a solid support, called “supported membrane” (Sackmann, 1996; Sackmann & Tanaka, 2021), which are functionalized with histidine-tagged human laminin E8 fragment. E8 fragment corresponds to the C-terminal integrin-binding region of laminin α_5_β1γ1.(Deutzmann *et al*, 1990; Miyazaki *et al*, 2012) In this study, we model the extracellular environments of PDA cells by functionalizing the surface with E8 fragment, because laminin α_5_β_1_γ_1_ is expressed in basement membranes of pancreatic ducts (Jiang *et al*, 2002). Supported membranes precisely functionalized with ligands offer several major advantages over solid substrates displaying immobilized ligands. Lipid membranes avoid non-specific adhesion of proteins and cells (Kaindl *et al*, 2012). Moreover, if one takes the area per lipid molecules in fluid phase *A*_lipid_ ≈ 0.7 nm^2^ (Lipowsky & Sackmann, 1995), the average intermolecular distance between laminin fragments on the membrane surface can be controlled at 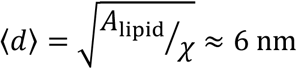 by doping the lipid anchors at a molar fraction of *𝓍* = 0.02 (Tanaka, 2019; Tanaka & Lanzer, 2022). **Figure 2b** presents the phase contrast images of IPMN-PDA and PanIN-PDA cells, which are distinct from those observed on plastic dishes (**Figure 1e**). Some IPMN-PDA cells exhibit pronounced axial stretching (indicated by arrows), while PanIN-PDA cells show no extensive elongation. Although the cell area showed no significant difference, the circularity index of IMPN-PDA cells, defined as 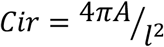, is significantly larger (*p* < 0.001) than that of PanIN-PDA cells (**Figure S2**), where *l* is cell contour length and *A* cell area. These data demonstrate that lamini-functionalized supported membranes can be used to discriminate the differential remodeling of adhesion contacts of PDA cells, which cannot be realized by conventional plastic dishes.

**Figure 2.**
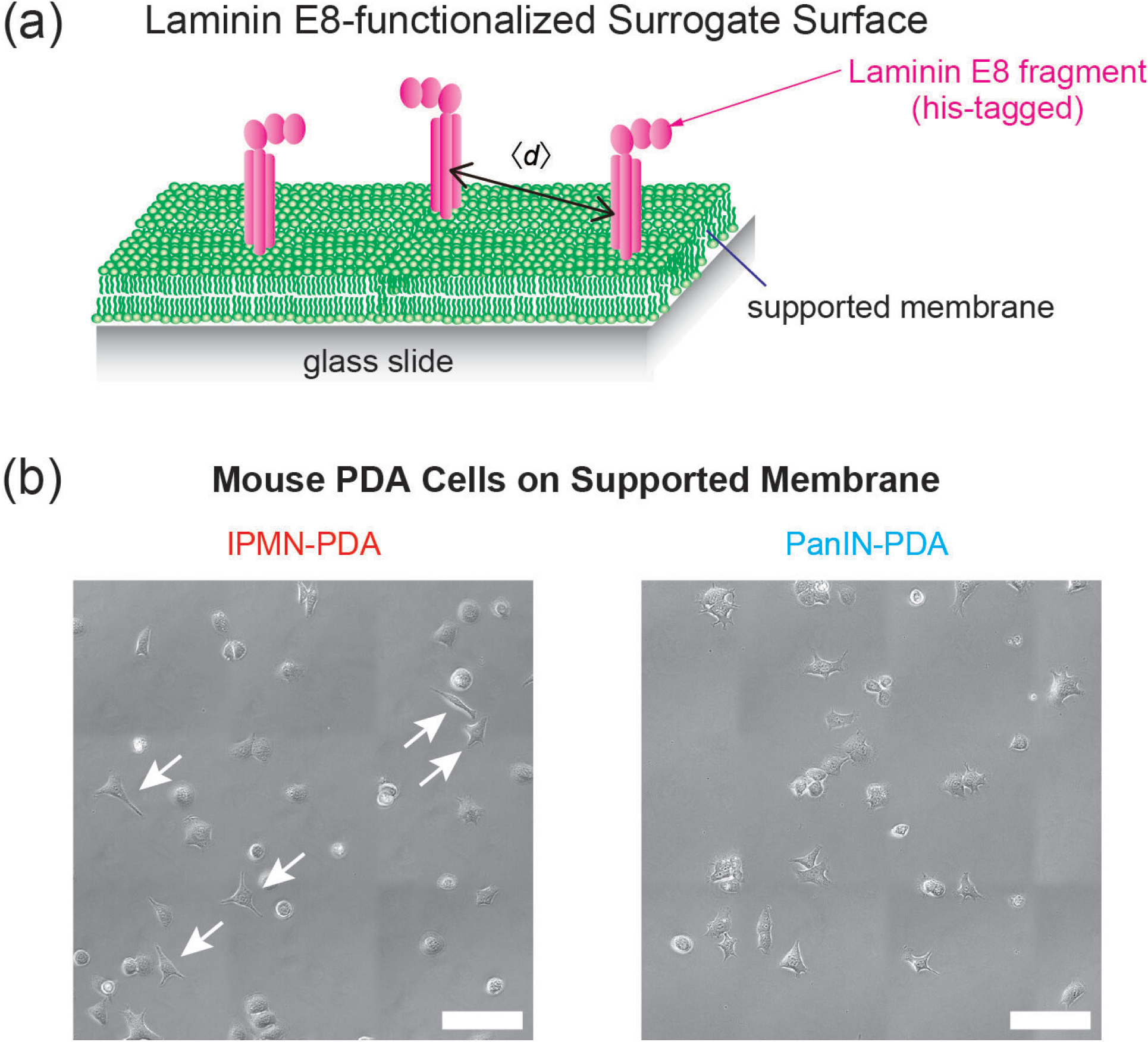
Laminin-functionalized supported membranes enables a clear discrimination of IPMN-PDA and PanIN-PDA. (a) Schematic illustration of a membrane-based surrogate surface displaying histidine-tagged Laminin E8 fragment. The average intermolecular distance between laminin fragments ⟨*d*⟩ was controlled to be 6 nm by the molar fraction of anchor lipids. (b) Phase contrast images of mouse IPMN-PDA and PanIN-PDA cells on supported membranes. Some IPMN-PDS cells show pronounced axial stretch, extending long filopodia (indicated by arrows). Scale bars: 50 µm.

### Membrane-based surrogate surfaces unravel distinct migration dynamics associated with chromatin remodeler Brg1

On supported membranes, PDA cells exhibit distinct dynamic phenotypes (**Movies S1 and S2**). Here, we performed live-cell imaging using refection interference contrast microscopy (RICM), which can detect the adhesion contact precisely in a label-free manner (Albersdörfer *et al*, 1997; Kaindl *et al*., 2012). **Figure 3a** shows the contour overlay of IPMN-PDA recorded over 500 min. The centroid position is defined from the cell contour, and the corresponding centroid trajectory is plotted in **Figure 3b**. In general, IPMN-PDA cells show pronounced axial stretch but little migration. The migration trajectories of IPMN-PDA cells (*N* = 20) are summarized in **Figure 3c**. Each line represents the trajectory of one cell and the starting point of each trajectory is shifted to the origin. Except for two cells (10%), the trajectories remain within 15 µm from the starting point, suggesting that the extensive axial stretch does not lead to migratory motion.

**Figure 3.**
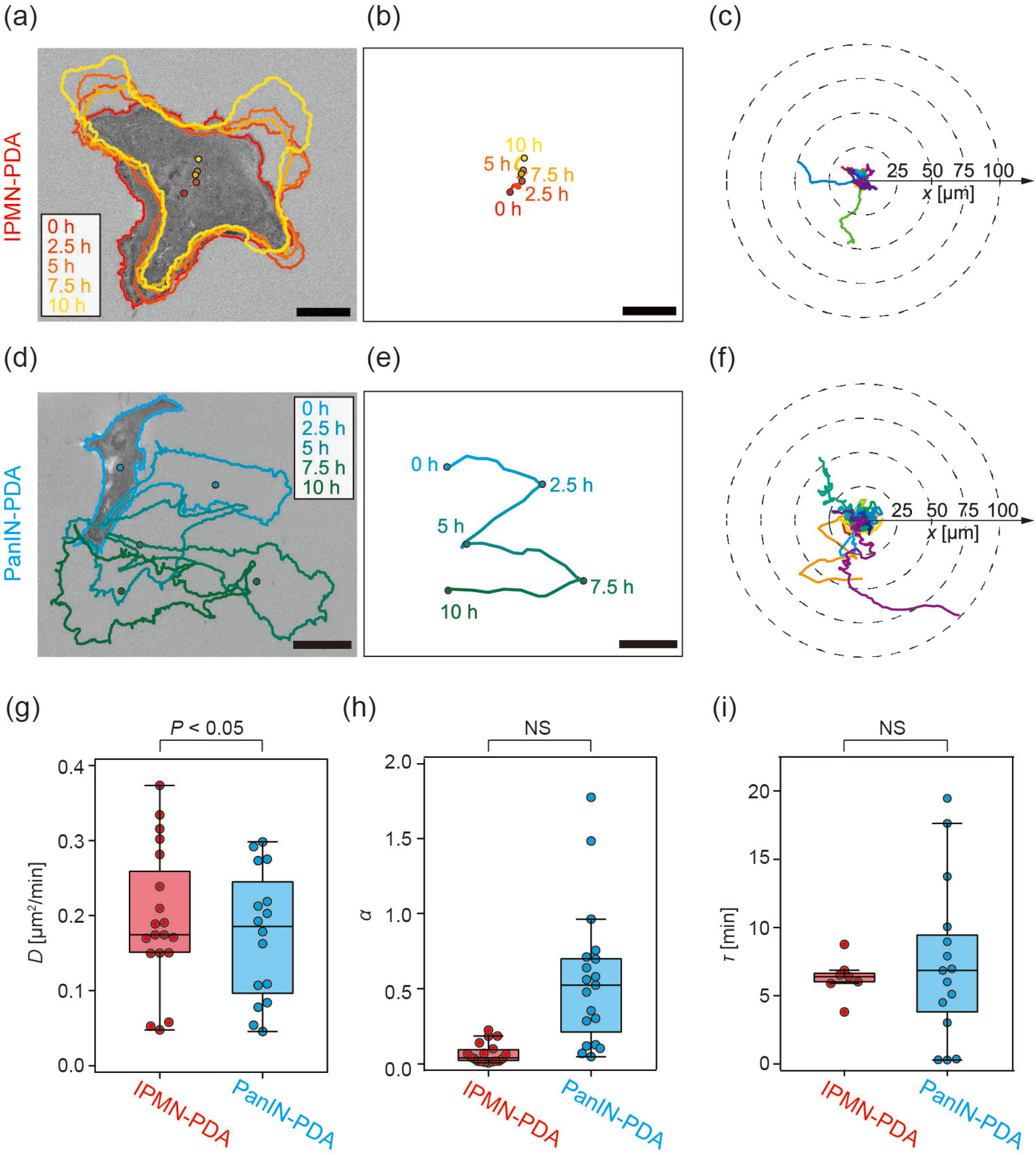
Distinct migrations of IPMN-PDA cells and PanIN-PDA cells. (a) (a) RICM images of IPMN-PDA single cell. The time course of cell contours over 10 h is indicated by color codes. (b) The trajectory of centroid at each time pointis defined from the contour extracted from a RICM snapshot. (c) Centroid trajectories of IPMN-PDA (*N* = 20, respectively) recorded over 10 h. The centroid position at *t* = 0 is defined as the origin for each cell. The corresponding datasets for PanIN-PDA are presented in panels (d)–(f). (g) Diffusion constant *D* and (h) power law exponent *α* of IPMN-PDA and PanIN-PDA calculated from MSD analysis. (i) Persistent time τ of IPMN-PDA and PanIN-PDA. Scale bars: 20 µm.

In contrast, PanIN-PDA cell undergoes dynamic deformation and migration. **Figures 3d** and **3e** represent the contour overlay and the centroid trajectory of PanIN-PDA cell. The migration trajectories of PanIN-PDA cells (*N* = 20) are plotted in **Figure 3f**. As a general trend, the trajectories of PanIN-PDA cells are more stretched compared to those of IMPN-PDA. From the trajectories, the mean squared displacement (MSD) profiles for individual cells are calculated (Eq. 1). From the double logarithmic plots of MSD as a function of lag-time Δ*t* (**Figure S3**), the diffusion coefficient *D* can be calculated. As presented in **Figure 3g**, IPMN-PDA exhibits the median diffusion coefficient of ⟨*D*_IPMN-PDA_⟩ = 0.014 μm^2^/min, which is almost one order of magnitude smaller than that of PanIN-PDA, ⟨*D*_PanIN-PD_ ⟩ = 0.11 μm^2^/min. The power law exponent *α* determined from the slope of double logarithmic plot, MSD *∝* (Δ*t*)^*α*^, is plotted in **Figure 3h**. The median exponent values of both PDA cells lay at around ⟨*α*⟩ ≈ 1, indicating that both PDA cells undergo random walks despite of a large difference in diffusion coefficients. This seems consistent with the experimental conditions, because sparsely distributed cells are not guided by any chemokine gradient. In both cell types, several cells show lower exponent values (*α* < 0.5), which can be attributed to the loss of polarity or pinning. The ballistics of motion is assessed by the persistent time of migrating direction τ, which is nothing but the correlation of migratory direction (Eq. 2, **Figure S4**) (Ohta *et al*., 2018). The calculated τ values are plotted in **Figure 3i**. PanIN-PDA cells exhibit a higher scatter, but the median levels for both cell types are comparable ⟨τ⟩ ≈ 6 min These data demonstrate that the migration of PDA is governed by random walk of sparsely distributed cells. Our data seem consistent with previous numerical simulations suggesting that ballistic motion only occurs at a high cell density (Vennettilli *et al*, 2022).

### Spatiotemporal analyses unravel modulation of dynamic deformation associated with Brg1

As presented in **Movies S1 and S2**, IPMN-PDA and PanIN-PDA exhibit distinct differences not only in migratory motion but also dynamic deformation. Here, we analyzed the deformation patterns of PDA cells to unravel how Brg1-associated signaling modulates active deformation (*N* = 20 for both PDAs). First, the radial distance between the centroid and periphery recorded at time point *t* is plotted in polar coordinate (**Figure 4a**), then the spatiotemporal patterns of deformation are calculated from the map of fluctuation amplitude *R*(θ, *t*) = *r*(θ, *t*) − ⟨*r*(θ, *t*)⟩_<sub>θ</sub>_ in real space (*C*_*RR*_(Δθ, Δ*t*)) and reciprocal space (Γ_m_) (**Figure 4b**).

**Figure 4.**
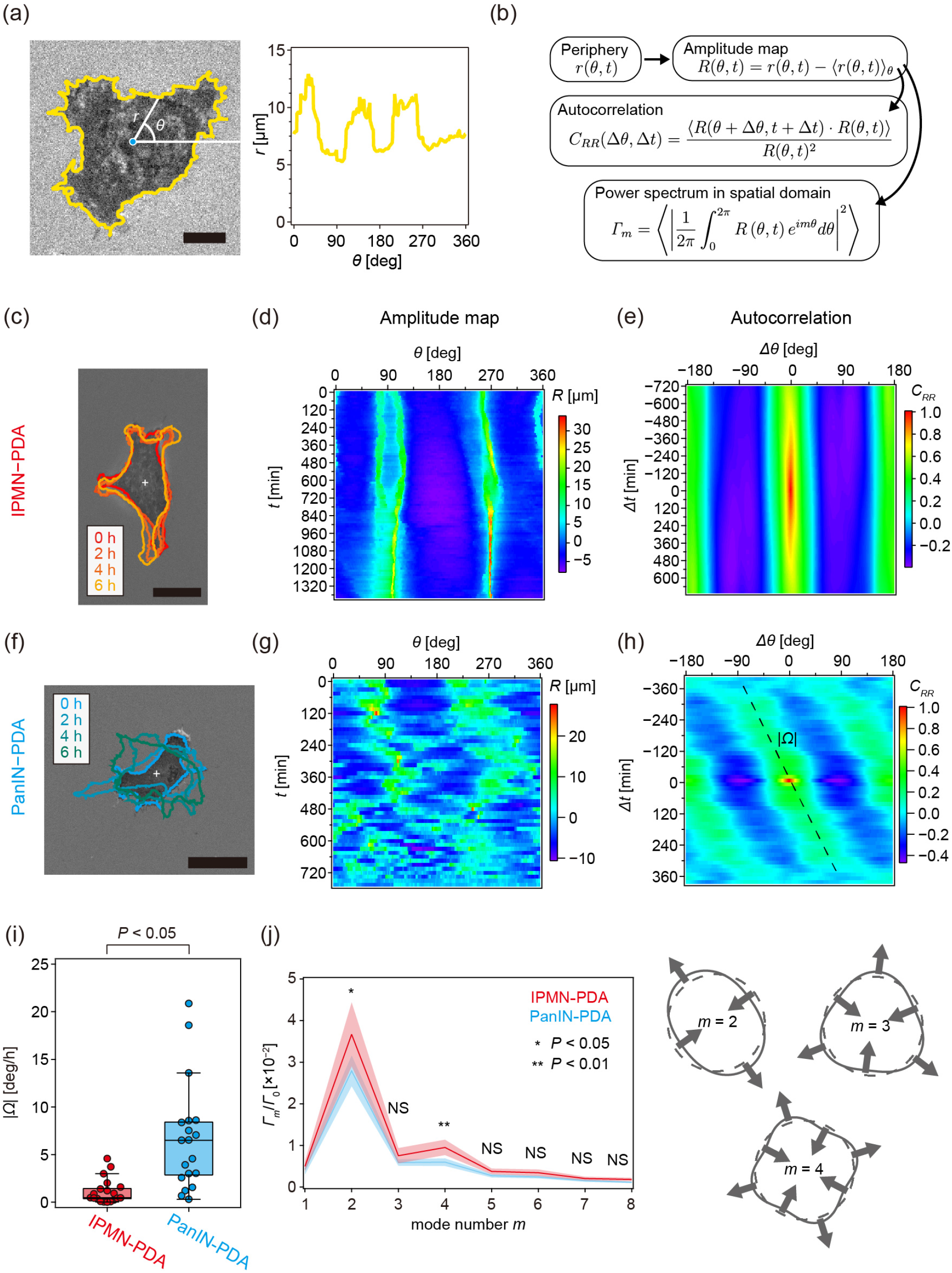
Deformation patterns of IPMN-PDA and PanIN-PDA in spatial dimension. (a) Left: An RICM image of a PanIN-PDA cell, whose contour is indicated by a blue line.Scale bar: 5 µm. Right: centroid-contour distance recorded at time point *t* plotted in polar coordinate, *r*(θ, *t*), yielding the fluctuation amplitude *R*(θ, *t*) = *r*(θ, *t*) − ⟨*r*(θ, *t*)⟩. (b) Flow of spatiotemporal analyses presented in the figure. The overlayed cell contours (c), amplitude map *R*(θ, *t*) (d), and correlation map *C*_*RR*_(Δθ, Δ*t*) (e) of IPMN-PDA. The corresponding datasets for PanIN-PDA are presented in panels (f)–(h). Broken line indicates the rotation. (i) Statistical comparison of |Ω| of IPMN-PDA and PanIN-PDA (*N* = 20, respectively). (j) Normalized power spectrum of cell deformation in spatial dimension Γ_m_, 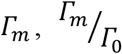. Γ_0_ is the squared average radius ⟨*r*(θ, *t*)^2^⟩_t_. The deformation in Fourier modes *m* = 2 ~ 4 are schematically illustrated.

**Figure 4c** represents the overlay of IPMN-PDA contours recorded over time. Here, the centroid position was fixed (indicated by a symbol) to highlight the shape fluctuation. **Figure 4d** shows the amplitude map (*R*(θ, *t*) map) of IPMN-PDA, exhibiting stable peaks at θ ≈ 90° and 270° that persist over *t*-axis. To extract characteristic deformation patterns hidden behind the random noise in real space, we calculated the autocorrelation function map *C*_*RR*_(Δθ, Δ*t*) (Eq. 3). **Figure 4e** presents the autocorrelation maps (*C*_*RR*_(Δθ, Δ*t*) map) of the IPMN-PDA. The two ridges at θ ≈ 90° and 270° lasting for Δ*t* ~ ± 600 min indicate that the cell is axially stretched in the same direction persistently. *R*(θ, *t*) maps and *C*_*RR*_(Δθ, Δ*t*) maps of other IPMN-PDA cells are presented in **Figures S5 and S6**, respectively.

In contrast, the contour overlays (**Figure 4f**) and the *R*(θ, *t*) map (**Figure 4g**) of PanIN-PDA seems noisy. Although two ridges at θ ≈ 90° and 270° in the *C*_*RR*_(Δθ, Δ*t*) map suggest an axial stretch, the ridges are less persistent than those of IPMN-PDA (**Figure 4h**). Moreover, the ridges are diagonally tilted with respect to the time axis (broken line), suggesting that the direction of elongation rotates over time. The tilt angle, 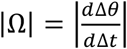, represents the angular velocity of an ellipse rotation: *R*(θ, *t*) ≈ *A* cos{2(θ − |Ω|*t*) − θ_0_}, where *A* and θ_0_ represent the amplitude and phase, respectively. The |ω| values of IPMN-PDA and PanIN-PDA are statistically compared in **Figure 4i**. The median level of IPMN-PDA is very low, ⟨|Ω|_IPMN-PDA_⟩ ≈ 0.5 deg/h, suggesting that IPMN-PDA hardly rotates. In contrast, the |Ω| values of PanIN-PDA cells show a markedly large scatter, taking the median value, ⟨|Ω|_PanIN-PDA_⟩ ≈ 6.5 deg/h. This indicates that PanIN-PDA rotates and changes the elongation direction. In fact, invasive cancer cells show irregular migration in order to move and spread fast (West *et al*, 2017). *R*(θ, *t*) maps and *C*_*RR*_(Δθ, Δ*t*) maps of other IPMN-PDA cells are presented in **Figures S7 and S8**, respectively.

The deformation of cells is an active, nonequilibrium process accompanied by energy consumption, such as bending of membranes, rearrangement of lipids and proteins, remodeling of cytoskeletons, and polarization and migration of cells (Burk *et al*, 2015; He *et al*, 2022; Lamas-Murua *et al*, 2018; Partin *et al*, 1989). Here, we calculated the power spectrum of cell deformation in spatial domain Γ_m_ to assess the energy consumed by PDA-cells via active deformation and migration (Burk *et al*., 2015; Lamas-Murua *et al*., 2018; Partin *et al*., 1989). *m* is the spatial mode of Fourier expansion series. To exclude the effect of cell size, we normalized Γ_m_by Γ_0_, which is the squared average radius ⟨*r*(θ, *t*)^2^⟩_t_(Eq. 4). In **Figure 4j**, the normalized power spectra 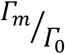 calculated for IPMN-PDA and PanIN-PDA are presented. Solid lines are mean values from *N* = 20 cells for each line. The 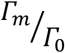 values of IPMN-PDA are larger than those of PanIN-PDA for all *m*,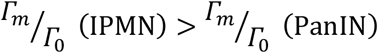, suggesting that the shape deformation of IPMN-PDA is larger than that of PanIN-PDA. The major peaks at *m* = 2 for both cell lines indicate that both IPMN-PDA and PanIN-PDA undergo predominantly axial deformation, as schematically illustrated in **Figure 4j**. Moreover, IPMN-PDA exhibit the larger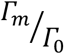 values at *m* = 4 than PanIN-PDA (**, *p* < 0.01). In contrast, the 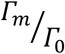 values at *m* = 3 are comparable. This suggests that IPMN-PDA cells spread in axial and biaxial directions but do not exhibit a clear front-rear asymmetry.

To further assess temporal dynamics of active cell deformation, the power spectrum in the temporal dimension 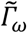 was calculated (Eq. 5). The flow of analyses in temporal dimension is schematically shown in **Figure 5a. Figure 5b** presents the normalized power spectra in frequency domain, 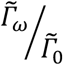, plotted in a double-logarithm plot.

**Figure 5.**
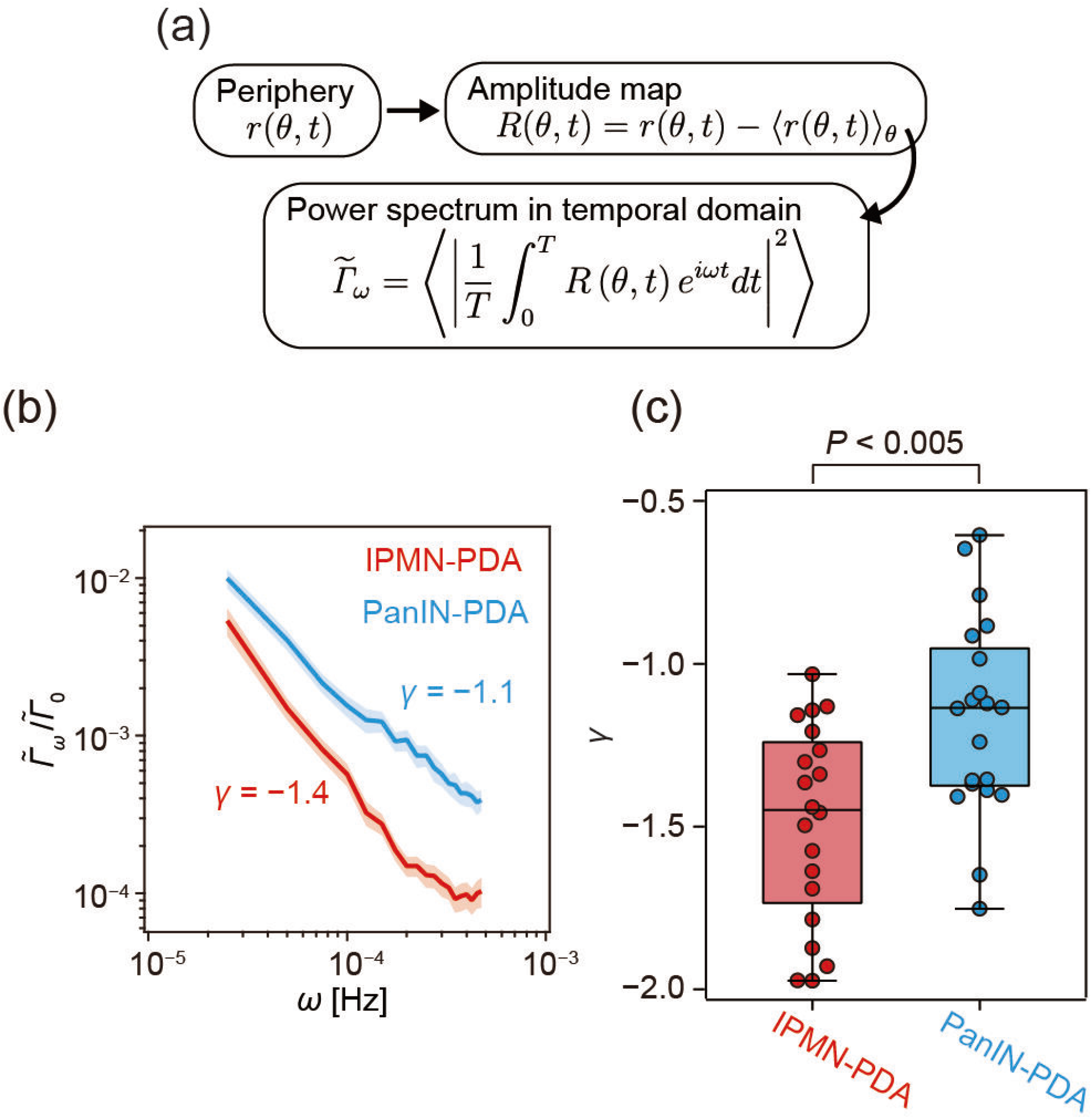
Deformation patterns of IPMN-PDA and PanIN-PDA in temporal dimension. (a) Flow of spatiotemporal analyses presented in the figure. (b) Power spectrum of cell deformation in temporal dimension 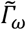 normalized by 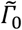 as a function of frequency ω, following 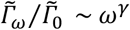. (c) Statistical comparison of power-law exponent *γ* values calculated from *N* = 20 cells for each line.

As shown in the figure, the temporal power spectrum density follows 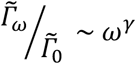, and the the power-low exponent γ reflects the rate of shape deformation. Of note, 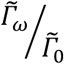the values of PanIN-PDA are larger than those of IPMN-PDA over the whole frequency range, indicating that the contour of PanIN-PDA cells fluctuates more dynamically than IPMN-PDA,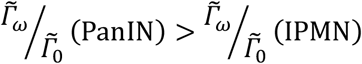. This is in contrast to the tendency observed in spatial dimension, 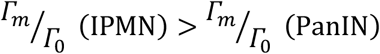. The statistical comparison of γ values is presented in **Figure 5c**. The median power-law exponent of PanIN-PDA, ⟨γ_PanIN-PDA_⟩ ≈ −1.1, is significantly larger than that of IPMN-PDA, ⟨γ_IPMN-PDA_⟩ ≈ −1.4 (*p* < 0.005). These data demonstrate that PanIN-PDA cells deform more actively and faster than IPMN-PDA.

Together, the quantitative analysis of motion and deformation of PDA cells highlighted the effects of chromatin remodeler Brg1 resulting in differential cellular dynamics both in space and in time. The slow and compact migration of IPMN-PDA can be attributed to a prominent bipolar elongation of IPMN-PDA with little front-rear asymmetry and slow shape relaxation. In contrast, the faster and more extended migration patterns of PanIN-PDA can be explained by the dynamic fluctuation and irregular rotation, accompanied by fast shape relaxation.

### Rac1 activity is associated with dynamic migration phenotypes and required for stable adhesion

To explore the molecular basis underlying the distinct dynamic behaviors of IPMN- and PanIN-derived PDA cells, we focused on the small GTPase Rac1, a key regulator of actin cytoskeleton remodeling and adhesion dynamics. To quantitatively assess Rac1 activity, the ratio of active, phosphorylated Rac1 (p-Rac1) to total Rac1 was determined by a pull-down assay (**Figure 6a**). As shown in **Figure 6b**, PanIN-PDA exhibited significantly higher levels of active Rac1 compared to IPMN-PDA (*p* < 0.05), indicating enhanced Rac1 signaling in the more dynamically migrating PanIN-PDA. Given the established role of Rac1 in lamellipodia formation and integrin-mediated adhesion, these results suggest that elevated Rac1 activity may contribute to the increased actin turnover and adhesion remodeling observed in PanIN-derived cells. This is consistent with the more dynamic deformation patterns and higher migration rates quantified in the analyses presented in the previous sections.

**Figure 6.** Rac1 activity in PanIN-PDA cells is associated with enhanced dynamics. (a) Rac1 pull-down assay of IPMN-PDA and PanIN-PDA. (b) PanIN-PDA exhibited significantly higher levels of active Rac1 compared to IPMN-PDA (*p* < 0.05). (c) RICM images of PanIN-PDA in the absence (control) and presence of 100 µM NSC23766 taken at *t* = 0 and 24 h. Scale bars: 20 µm. (d) Adhesion to laminin-functionalized surfaces decreases in a concentration depending manner, indicating that Rac1 activity is required for maintaining stable adhesion. Error bars indicate 95% confidence intervals calculated using the Wilson score method (n > 45 for each condition).

Previously, we employed dedifferentiated *Kras*^G12D^ and *Brg1*^f/f^ acinar cell explants and demonstrated that Brg1 binds to the Sox9 promoter region, and this Brg1/Sox9 axis plays critical roles in acinar cell-derived tumorigenesis in pancreas (Tsuda *et al*, 2018). From the microarray data in (Tsuda *et al*., 2018), we found that guanine nucleotide exchange factor (GEF) genes involved in Rho GTPase regulation, *Trio, Tiam2, Dock1, Dock7, Arhgef39, Arhgef26*(*SGEF*), and *Arhgef2*(*GEF-H1*) are downregulated in *Brg1*^f/f^ acinar cell explants (**Table S1**) (Tsuda *et al*., 2018). Together, these data suggests that Brg1-dependent processes are associated with the dedifferentiation of acinar cells into a PanIN state, a process that requires and activates Rac1 pathway.

To further examine the functional role of Rac1in this system, we performed pharmacological inhibition experiments using the Rac1-specific inhibitor NSC23766.

**Figure 6c** shows the RICM images of PanIN-PDA at *t* = 0 h and 24 h in the absence (DMSO) and presence of 100 µM NSC23766. Adhesion contacts of the cells treated with 100 µM NSC23766 decreased, and some cells detached after 24 h. At a low concentration (25 μM), no detectable changes in migration trajectories or deformation dynamics were observed compared to untreated cells. In contrast, at higher concentrations (50–100 μM), cells progressively lost adhesion to the laminin-functionalized supported membrane. Quantification of the fraction of attached cells as a function of NSC23766 concentration confirmed a marked reduction in adhesion stability under these conditions (**Figure 6d**). These observations indicate that Rac1 activity is required for maintaining stable adhesion on laminin-functionalized membranes. However, the loss of adhesion at higher inhibitor concentrations precluded a direct assessment of Rac1-dependent changes in migration dynamics independent of adhesion. Together, these results support a model in which elevated Rac1 activity in PanIN-PDA cells is associated with enhanced cytoskeletal dynamics and adhesion remodeling, contributing to their distinct migratory behavior.

## Discussion

In this study, we establish a quantitative framework that links epigenetic state to the spatiotemporal dynamics of cell migration by integrating biomimetic surfaces with quantitative analysis of cell motion and deformation. Our results show that cancer cells derived from distinct precursor lesions (IPMN and PanIN) can unravel differential physical phenotypes despite being indistinguishable by conventional morphological criteria, underscoring the importance of quantitative analyses of spatiotemporal dynamics in resolving functional heterogeneity. By integrating trajectory analysis with Fourier-based characterization of deformation, we identify two distinct modes of cellular behavior: a of slow, persistent deformation with limited net displacement and a of rapid, fluctuating remodeling associated with enhanced migration. These modes are not captured by static morphological descriptors on plastic dishes but emerge clearly in the spatiotemporal domain on functionalized supported membranes, suggesting that dynamic physical features provide a sensitive readout of underlying regulatory states. Our molecular analyses indicate that Rac1 activity is associated with these distinct dynamic patterns downstream of the chromatin remodeler Brg1. PanIN-PDA cells exhibit elevated levels of active Rac1, consistent with their increased actin turnover, adhesion remodeling, and enhanced migration. These findings support a model in which Brg1-dependent regulation modulates Rac1 activity, thereby influencing cytoskeletal dynamics and cell behavior.

To further probe the functional role of Rac1, we performed pharmacological inhibition experiments using NSC23766. While low concentrations had little effect on migration dynamics, higher concentrations led to a progressive loss of cell adhesion to the laminin-functionalized membrane, ultimately resulting in cell detachment. These observations indicate that Rac1 activity is required for maintaining stable adhesion under the present experimental conditions. The loss of adhesion at higher inhibitor concentrations prevented a direct assessment of Rac1-dependent migration dynamics independent of adhesion. Nevertheless, this behavior is consistent with the established role of Rac1 in coordinating actin polymerization and integrin-mediated adhesion (Sackmann & Tanaka, 2021). Together, these results suggest that elevated Rac1 activity in PanIN-derived cells promotes both cytoskeletal remodeling and adhesion dynamics, enabling the observed migratory phenotype.

The experimental approach introduced here provides a platform for probing dynamic cellular phenotypes. Supported lipid membranes functionalized with defined ligands offer a biomimetic interface that minimizes nonspecific interactions while allowing controlled presentation of adhesion molecules (Balta *et al*, 2019; Fröhlich *et al*, 2021; Kaindl *et al*., 2012). Combined with label-free interferometric imaging, this system enables long-term, quantitative measurements of cell behavior without perturbing cellular function. Importantly, the analytical framework based on spatiotemporal descriptors is broadly applicable and can be extended to other cell types and microenvironments.

Here, we report that epigenetic regulation encodes not only transcriptional programs but also dynamic physical phenotypes that may underlie invasive potential. From a broader perspective, our data suggest that epigenetic regulation may encode not only gene expression programs but also physical phenotypes that manifest in cell dynamics. This perspective provides a conceptual bridge between molecular biology and biophysics and highlights the potential of dynamic cellular features as quantitative biomarkers (Tanaka *et al*, 2023). Such descriptors may complement existing approaches by providing functional information about invasive potential, particularly in cancers where static morphological assessment is insufficient.

Several limitations of the present study should be noted. First, the analysis is based on *in vitro* systems that dissect cell-intrinsic dynamics from the complexity of tissue environments. Second, although Rac1 inhibition experiments demonstrate a requirement for Rac1 in maintaining adhesion, more targeted perturbations will be necessary to dissect its specific contributions to migration and deformation dynamics. Finally, for evaluating its relevance to disease progression, it is important to connect this framework to *in vivo* contexts.

## Conclusion

In this study, we combined biomimetic interfaces with quantitative analysis of cell dynamics to investigate how epigenetic regulation shapes cancer cell behavior. We show that pancreatic cancer cells derived from distinct precursor lesions exhibit markedly different migration and deformation dynamics despite almost indistinguishable morphology. These differences are associated with Rac1 activity downstream of the chromatin remodeler Brg1 and are reflected in distinct modes of cytoskeletal and adhesion dynamics.

Our results highlight the importance of dynamic cellular phenotypes as functional readouts of molecular regulation. By linking epigenetic state to measurable spatiotemporal features of cell behavior, this work provides a framework for connecting molecular mechanisms to emergent physical properties. Such approaches are broadly applicable and may contribute to the development of quantitative descriptors that capture invasive potential and functional heterogeneity in cancer.

## Methods

### Materials

1,2-dioleoyl-sn-glycero-3-phosphocholine (DOPC) and 1,2-dioleoyl-sn-glycero-3-[(N- (5-amino-1-carboxypentyl)iminodiacetic acid)succinyl] (nickel salt) (DGS-NTA (Ni^2+^)) were purchased from Avanti Polar Lipids (Alabaster, AL, United States). The recombinant of laminin E8 (iMatrix-511) was purchased from Nippi (Tokyo, Japan). Unless stated otherwise, all chemicals were purchased from Nacalai Tesque (Kyoto, Japan) and used without further purification.

### Preparation of membrane-based surrogate surface

Glass substrates (25 × 75 mm^2^) were cleaned following previous reports (Hillebrandt & Tanaka, 2001; Kern, 1970) and sealed with bottomless plastic chambers (Ibidi, Martinsried, Germany) (Burk *et al*., 2015; Kaindl *et al*., 2012). Supported membranes were prepared by fusion of DOPC vesicles incorporating 2 mol% DGS-NTA in 150 mM NaCl buffered with 10 mM HEPES (pH7.4). After removal of excess vesicles, the supported membrane was incubated with 1 mM NiCl_2_ for 30 min, rinsed with the buffer, and incubated with laminin E8 solution (10 μg/ml) for 60 min. After removing non-bound proteins, the chambers were equilibrated at 37 °C (Burk *et al*., 2015; Tanaka *et al*., 2023).

### Cell culture

PanIN-PDA and IPMN-PDA cell lines were established as previously reported (von Figura *et al*., 2014). The culture medium was composed of Dulbecco’s modified eagle medium (DMEM) (#11995-065, Gibco), heat-inactivated 10% fetal bovine serum (FBS) (#10270106, Gibco), and 1% penicillin-streptomycin (PS, P0781, Sigma-Aldrich). The cells were harvested using 0.25% Trypsin-EDTA solution (#T4049, Sigma-Aldrich) and passaged every 2 or 3 days. For live-cell imaging, cells were suspended in FBS(−) medium and 1 × 10^4^ cells were seeded into a well with the area of 1.0 cm^2^ which was filled with FBS(−) medium. Cells were allowed to adhere for 1 h prior to the live imaging.

### Live-cell imaging

Live-cell imaging was performed using reflection interference contrast microscopy (RICM) on an Axiovert.Z1 inverted microscopy (Carl Zeiss, Oberkochen, Germany) equipped with an oil-immersion objective lens (NA 1.25, 63 ×, PH3) and an Orca-Flash4.0 camera (Hamamatsu Photonics, Hamamatsu, Japan. Images were acquired every 15 min over 48 h, and the exposure time was kept constant (100 ms). The contour of adhesion zone was extracted by image binarization using a custom-made algorithm in ImageJ Fiji (NIH, USA).

### Spatiotemporal analysis

The analysis of spatiotemporal dynamics was performed using custom-made algorithms in IGOR Pro (WaveMetrics, Portland, OR, USA). To characterize the migratory behavior of cells, the centroid position of the adhesion zone 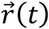 was used to calculate MSD as a function of lag time Δ*t* as

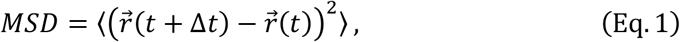

where 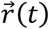 is the centroid position of the adhesion zone at time *t* and the bracket ⟨ ⟩ indicates to take the average over *t*. The persistence time of trajectory τ was calculated as the exponential decay of the migrating direction:

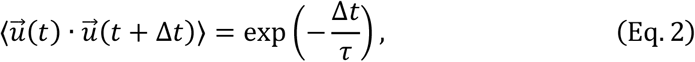

where 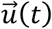 represents the unit tangential vector of the trajectory at time *t*.

The spatiotemporal dynamics of the adhesion zone were quantitatively analyzed via the fluctuation amplitude of centroid-contour distance recorded over time, *R*(θ, *t*) = *r*(θ, *t*) − ⟨*r*(θ, *t*)⟩_0_. To extract characteristic patterns, the autocorrelation *C*_*RR*_(Δθ, Δ*t*) was determined as

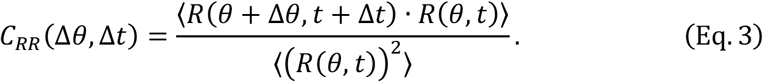

The angular velocity of an ellipse rotation |ω| was calculated from the 2D-FFT (Fast Fourier Transformation) of the *R*(θ, *t*) map. The power spectrum of cell deformation in spatial domain Γ_m_ and temporal domain 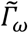 were determined from the 1D-FFT of the *R*(θ, *t*) map as

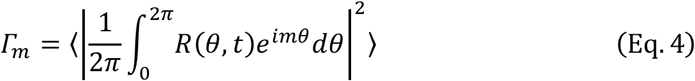

and

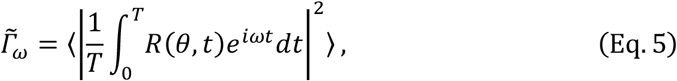

where *m, ω*, and *T* are the mode number, rotational frequency, and the duration of observation, respectively.

### Protein Analysis

The ratio of activated Rac1 and total Rac1 expressed in IPMN-PDA and PanIN-PDA was quantified by Rac1 Pull-down Activation Assay Biochem Kit (#BK035, Cytoskeleton). The pulled-down p-Rac1 was resolved on SDS-PAGE and analyzed by Western blotting using Clarity Western ECL Substrate (#1705060, Bio-Rad).

### Statistical analysis

Statistical analysis was performed in IGOR Pro. All box plots presented in this study present the median value as a solid line. The boxes correspond to the interquartile ranges (IQR), and the whiskers extend to the most extreme values within 1.5 × *IQR* from first (*Q*_l_) and third (*Q*_3_) quartiles, where *IQR* = *Q*_3_ − *Q*_l_. Data points beyond 3.0 × *IQR* from first and third quartiles were regarded as outliers. Comparisons between two groups were performed using Student’s *t* test. The *p* values < 0.05 were considered as significant difference.

## Data Availability Statement

The authors declare that the main data supporting the results in this study are available within the paper and its Supplementary Information. The raw and analyzed datasets generated during the study are available for research purposes from the corresponding authors (M.T. and H.S.) on reasonable request.

## Author Contributions

**Akihisa Yamamoto**: Data Acquisition; Analysis; Writing–original draft. **Akihisa Fukuda**: Resources; Conceptualization; Supervision; Methodology; Writing–review and editing. **Yuichi Fukunaga**: Resources; Analysis. **Kentaro Hayashi**: Analysis. **Hiroshi Seno**: Conceptualization; Supervision; Fund Acquisition; Methodology; Writing–review and editing, **Motomu Tanaka**: Conceptualization; Supervision; Analysis; Fund Acquisition; Writing–original draft, review and editing.

## Disclosure and competing interests statement

The authors declare no conflict of interests.

## Acknowledgement

The authors thank Satoko Hinatsu and Moritz Tremmel for the technical assistance. This study was supported by JSPS KAKENHI (JP20H00661 to M.T. and H.S., JP24H00796 to M.T.) and the German Science Foundation (Germany’s Excellence Strategy— 2082/1—390761711 to M.T.). M.T. also thanks the Nakatani Foundation for its support. The authors thank German-Japanese HeKKSaGOn Alliance for support.

## References

Albersdörfer A, Feder T, Sackmann E (1997) Adhesion-induced domain formation by interplay of long-range repulsion and short-range attraction force: a model membrane study. Biophysical journal 73: 245–257

Balta GSG, Monzel C, Kleber S, Beaudouin J, Balta E, Kaindl T, Chen S, Gao L, Thiemann M, Wirtz CR et al (2019) 3D cellular architecture modulates tyrosine kinase activity, thereby switching CD95-mediated apoptosis to survival. Cell reports 29: 2295-2306.e2296

Basturk O, Hong S, Wood L, Adsay N, Albores-Saavedra J, Biankin A, Brosens L, Fukushima N, Goggins M, Hruban R (2015) Baltimore Consensus Meeting. A revised classification system and recommendations from the Baltimore consensus meeting for neoplastic precursor lesions in the pancreas. Am J Surg Pathol 39: 1730–1741

Burk AS, Monzel C, Yoshikawa HY, Wuchter P, Saffrich R, Eckstein V, Tanaka M, Ho AD (2015) Quantifying adhesion mechanisms and dynamics of human hematopoietic stem and progenitor cells. Scientific reports 5: 9370

Deutzmann R, Aumailley M, Wiedemann H, Pysny W, Timpl R, Edgar D (1990) Cell adhesion, spreading and neurite stimulation by laminin fragment E8 depends on maintenance of secondary and tertiary structure in its rod and globular domain. European journal of biochemistry 191: 513–522

Fröhlich B, Dasanna AK, Lansche C, Czajor J, Sanchez CP, Cyrklaff M, Yamamoto A, Craig A, Schwarz US, Lanzer M et al (2021) Functionalized supported membranes for quantifying adhesion of P. falciparum-infected erythrocytes. Biophysical Journal 120: 3315–3328

Fukuda A (2015) Molecular mechanism of intraductal papillary mucinous neoplasm and intraductal papillary mucinous neoplasm-derived pancreatic ductal adenocarcinoma. Journal of Hepato-Biliary-Pancreatic Sciences 22: 519–523

Habib JR, Rompen IF, Javed AA, Grewal M, Kinny-Köster B, Andel PCM, Hewitt DB, Sacks GD, Besselink MG, van Santvoort HC et al (2024) Outcomes in intraductal papillary mucinous neoplasm-derived pancreatic cancer differ from PanIN-derived pancreatic cancer. Journal of Gastroenterology and Hepatology 39: 2360–2366

He L, Arnold C, Thoma J, Rohde C, Kholmatov M, Garg S, Hsiao CC, Viol L, Zhang K, Sun R et al (2022) CDK7/12/13 inhibition targets an oscillating leukemia stem cell network and synergizes with venetoclax in acute myeloid leukemia. EMBO Molecular Medicine 14: EMMM202114990

Hillebrandt H, Tanaka M (2001) Electrochemical characterization of self-assembled alkylsiloxane monolayers on indium− tin oxide (ITO) semiconductor electrodes. The Journal of Physical Chemistry B 105: 4270–4276

Jiang F-X, Naselli G, Harrison LC (2002) Distinct distribution of laminin and its integrin receptors in the pancreas. Journal of Histochemistry & Cytochemistry 50: 1625–1632

Kaindl T, Rieger H, Kaschel L-M, Engel U, Schmaus A, Sleeman J, Tanaka M (2012) Spatio-temporal patterns of pancreatic cancer cells expressing CD44 isoforms on supported membranes displaying hyaluronic acid oligomers arrays.

Kern W (1970) Cleaning solution based on hydrogen peroxide for use in silicon semiconductor technology. RCA review 31: 187–206

Kiemen AL, Dequiedt L, Shen Y, Zhu Y, Matos-Romero V, Forjaz A, Campbell K, Dhana W, Cornish T, Braxton AM et al (2024) PanIN or IPMN? Redefining Lesion Size in 3 Dimensions. The American Journal of Surgical Pathology 48: 839–845

Lamas-Murua M, Stolp B, Kaw S, Thoma J, Tsopoulidis N, Trautz B, Ambiel I, Reif T, Arora S, Imle A (2018) HIV-1 Nef disrupts CD4+ T lymphocyte polarity, extravasation, and homing to lymph nodes via its Nef-associated kinase complex interface. The Journal of Immunology 201: 2731–2743

Lipowsky R, Sackmann E (1995) Structure and dynamics of membranes: I. from cells to vesicles/II. generic and specific interactions. Elsevier

Miyazaki T, Futaki S, Suemori H, Taniguchi Y, Yamada M, Kawasaki M, Hayashi M, Kumagai H, Nakatsuji N, Sekiguchi K (2012) Laminin E8 fragments support efficient adhesion and expansion of dissociated human pluripotent stem cells. Nature communications 3: 1236

Ohta T, Monzel C, Becker AS, Ho AD, Tanaka M (2018) Simple physical model unravels influences of chemokine on shape deformation and migration of human hematopoietic stem cells. Scientific reports 8: 10630

Partin AW, Schoeniger JS, Mohler JL, Coffey DS (1989) Fourier analysis of cell motility: correlation of motility with metastatic potential. Proceedings of the National Academy of Sciences 86: 1254–1258

Pedro BA, Wood LD (2025) Early neoplastic lesions of the pancreas: initiation, progression, and opportunities for precancer interception. The Journal of Clinical Investigation 135

Sackmann E (1996) Supported membranes: scientific and practical applications. Science 271: 43–48

Sackmann E, Tanaka M (2021) Critical role of lipid membranes in polarization and migration of cells: a biophysical view. Biophysical Reviews 13: 123–138

Tanaka M (2019) In vitro dynamic phenotyping for testing novel mobilizing agents. In: Stem Cell Mobilization: Methods and Protocols, pp. 11–27. Springer:

Tanaka M, Lanzer M (2022) Receptor-functionalized lipid membranes as biomimetic surfaces for adhesion of Plasmodium falciparum-infected erythrocytes. In: Malaria Immunology: Targeting the Surface of Infected Erythrocytes, pp. 601–613. Springer:

Tanaka M, Thoma J, Poisa-Beiro L, Wuchter P, Eckstein V, Dietrich S, Pabst C, Müller-Tidow C, Ohta T, Ho AD (2023) Physical biomarkers for human hematopoietic stem and progenitor cells. Cells & Development 174: 203845

Tsuda M, Fukuda A, Roy N, Hiramatsu Y, Leonhardt L, Kakiuchi N, Hoyer K, Ogawa S, Goto N, Ikuta K et al (2018) The BRG1/SOX9 axis is critical for acinar cell–derived pancreatic tumorigenesis. The Journal of Clinical Investigation 128: 3475–3489

Vennettilli M, González L, Hilgert N, Mugler A (2022) Autologous chemotaxis at high cell density. Physical Review E 106: 024413

von Figura G, Fukuda A, Roy N, Liku ME, Morris Iv JP, Kim GE, Russ HA, Firpo MA, Mulvihill SJ, Dawson DW et al (2014) The chromatin regulator Brg1 suppresses formation of intraductal papillary mucinous neoplasm and pancreatic ductal adenocarcinoma. Nature Cell Biology 16: 255–267

West A-KV, Wullkopf L, Christensen A, Leijnse N, Tarp JM, Mathiesen J, Erler JT, Oddershede LB (2017) Dynamics of cancerous tissue correlates with invasiveness. Scientific Reports 7: 43800

Yu W, Zhou D, Meng F, Wang J, Wang B, Qiang J, Shen L, Wang M, Fang H (2025) The global, regional burden of pancreatic cancer and its attributable risk factors from 1990 to 2021. BMC Cancer 25: 186

